# Genomic analyses reveal FoxG as an upstream regulator of *wnt1* required for posterior identity specification in planarians

**DOI:** 10.1101/2020.12.08.416008

**Authors:** E. Pascual-Carreras, M. Marín-Barba, S. Castillo-Lara, P. Coronel-Córdoba, M.S. Magri, G.N. Wheeler, J.F. Abril, J.L. Gomez-Skarmeta, E. Saló, T. Adell

## Abstract

Embryonic specification of the first body axis requires the formation of an Organizer, a group of cells with the ability to instruct fates in the surrounding tissue. The existence of organizing regions in adults, i.e. during regeneration, which also requires patterning of new tissues, remains unstudied. To that aim, we study regeneration in planarians, flatworms that can regenerate any missing structure, even the head, in a few days. In planarians, as described in embryonic models, the cWNT pathway specifies the anterior-posterior axis. During the first 12-24h after amputation both *wnt1* and *notum* (a Wnt inhibitor) are expressed in any wound, but 48 hours later they become restricted to posterior or anterior facing wounds, forming the anterior and the posterior organizers, respectively. In this study we undertook a genomic approach to further understand the mechanism that triggers the early expression of *wnt1* and the specification of the posterior identity. Through ATAC-sequencing and CHIPmentation techniques we uncovered Cis-Regulatory Elements of *Schmidtea mediterranea* genome and analyzed them in *notum* and *wnt1* (RNAi) animals. The result shows that already at 12 hours after amputation the chromatin structure of the wounds has changed its conformation according to the polarity of the pre-existing tissue. Analysing the DNA binding motives present in the proximal regulatory regions of genes down-regulated after *wnt1* (RNAi) we found a few genes containing a TCF binding site, which include posterior Homeobox genes and chromatin remodelling proteins, suggesting that those are direct targets of the cWNT pathway and the responsible to trigger the expression of the posterior effectors. Furthermore, we have identified FoxG as an up-stream regulator of *wnt1* transcription, probably though binding to an enhancer found in its first intron. Silencing of *foxG* inhibits the early phase of *wnt1* expression and phenocopies the *wnt1* (RNAi) phenotype, indicating its early role in specifying posterior *versus* anterior identity. Moreover, we have created a new open platform to interpret all transcriptomic and genomic results obtained (https://compgen.bio.ub.edu/PlanNET/planexp).

## Introduction

During embryonic development specification of the body axis is one of the earliest events, since it creates a coordinate system to which refer when building all organs and tissues. Specification of the first body axis requires the formation of an Organizing Center or Organizer, which refers to a group of cells with the ability to instruct fates and morphogenesis in surrounding cells, giving rise to specific organs and tissues (1–3). Spemann and Mangold were the first to demonstrate that the dorsal lip of a newt early gastrula had the ability to generate a fully patterned secondary axis when grafted to the opposite site (4–8). The homologous organizer is found during gastrulation of all vertebrates, receiving different names, as the Hensen’s node in birds (9), dorsal shield in fish embryos (10). The difference between Organizers and Organizing centers is commonly attributed to their ability to pattern a whole body axis or just an organ or tissue, respectively. Organizing centers have been identified in several stages of development, for instances in the limb bud of tetrapods or the isthmic organizer at the midbrain–hindbrain boundary (1,11,12). Although organizers are commonly studied in embryos, the very first experiment that demonstrated the existence of an organizer was in fact performed in adult hydras by Ethel Browne in 1909. Ethel Browne transplanted non-pigmented head tissue into the body column of a pigmented hydra and observed the induction of a secondary axis that was composed by the cells of the host (13). More than a century later the existence of organizing regions in adults, i.e. during regeneration, which also requires re-patterning of tissues, is not well studied. To that aim, we study the process of regeneration of planarians, flatworms which can regenerate any missing structure, even the head, in a few days. Thus, they are whole body-regenerating animals, which need re-patterning the body axes to regenerate the proper missing structures according to the pre-existent polarity.

Planarians plasticity is based on the presence of a population of adult pluripotent stem cells (called neoblast) (14,15), together with the continuous activation of the signalling pathways that instruct the fate of these stem cells and their progeny. Several studies demonstrate that the muscle layer that surrounds the planarian body is the source of the so-called Positional Control Genes (PCGs), which are secreted factors that confer axial identity to the rest of the cells (16–19). A subset of these muscular cells located in the most anterior (tip of the head) and the most posterior (tip of the tail) ends of the planarian body act as Organizers. The anterior tip expresses *notum* (a secreted cWNT pathway inhibitor) and the posterior tip secretes *wnt1* (a cWNT pathway activator), and inhibition of those genes produces a shift in the polarity, originating two-tailed or two-headed planarians after silencing *notum* or *wnt1*, respectively (20–22). Thus, in planarians, as described in several embryonic models, the cWNT pathway specifies the anterior-posterior axis (23–28). Importantly, during the first 12-24 hours of regeneration (hR) both *notum* and *wnt1* are expressed in muscle cells of any wound, and it is not after 36 hR that they are restricted to the tip of the anterior or posterior facing wounds, respectively (20,22,29), forming the anterior and the posterior organizing regions. It is known that the late localized expression of *notum* and *wnt1* depends on the proliferation of stem cells, and that it requires the expression of some transcription factors, such as *foxD, zicA, prep* or *pbx* for anterior tips (30–34) and *islet, pitx* and *teashirt* (*tsh*) for posterior tips (35–38). However, the triggering of the early expression of *notum* and *wnt1*, which does not depend on stem cell proliferation, is not understood, as well as the molecular mechanism that restricts each factor to its corresponding pole and the following molecular events that take place to finally regenerate the missing structure.

In this study we undertook a genomic approach to analyse the formation process of the posterior Organizer during planarian regeneration. Through ATAC-sequencing and CHIPmentation techniques we uncovered Cis-Regulatory Elements (CREs) of *Schmidtea mediterranea* genome (39) and we analysed their accessibility in wild type (wt), *notum* and *wnt1* (RNAi) regenerating wounds. Our results show that already at 12 hR, anterior wounds of *notum* (RNAi) animals resemble wt posterior wounds, and posterior wounds of *wnt1* (RNAi) animals resemble wt anterior wounds. Thus, during the first hours after amputation, before the expression of any anterior or posterior marker, the chromatin structure of the wounds has already changed its conformation according to the polarity of the pre-existing tissue. Analysing the DNA binding motifs upstream of genes down-regulated after *wnt1* (RNAi) we found a few genes containing a TCF binding site, suggesting that those are the genes directly regulated by the cWNT pathway (Wnt1-βcatenin1) and responsible to trigger the posterior program. Finally, we identified a putative enhancer located in the first intron of *wnt1* containing a FoxG binding site. Silencing of *foxG* inhibits the early and the late phase of *wnt1* expression, but not *notum*, and phenocopies the wnt1 RNAi phenotype. This result suggests that FoxG directly regulates the early expression of *wnt1* in any wound and is a key factor in triggering the formation of the posterior organizing and thus specifying posterior *versus* anterior identity.

Finally, we have created a new open platform to query and interpret all transcriptomic and genomic results obtained (https://compgen.bio.ub.edu/PlanNET/planexp). An additional web-tool has been developed in order to search for transcription factor binding sites and to explore the newly predicted regulatory elements (https://compgen.bio.ub.edu/PlanNET/tf_tools).

## Results

### 12 hours after amputation the chromatin structure of cells in the wound has changed according to the polarity of the pre-existing tissue

To identify cis-regulatory elements (CREs) that after amputation could specify anterior or posterior identity, we performed ATAC-sequencing and CHIPmentation of anterior and posterior wounds 12 hours after post-pharyngeal amputation of *Schmidtea mediterranea* (Figure 1A). At this regeneration time point the early expression of the cWNT elements *notum* and *wnt1* is first detected, although it is still not polarized (20,22,29). The comparison of the results obtained when analysing anterior versus posterior wounds allowed us to identify ATAC-seq peaks corresponding to accessible chromatin regions (ACRs) specific for each pole (DiffBind, FDR<0.05, fc>2). We found 611 specific anterior ACRs and 2484 specific ACRs of posterior (Figure 1, Table S1). Comparing those ACRs with ChIPmentation of samples also corresponding to 12 hR anterior and posterior wounds using the H3K27ac antibody, which allows the identification of active enhancers (40,41), we were able to identify a list of 555 anterior putative active enhancers and 1869 posterior putative active enhancers (Figure 1 and Table S1).

**Figure 1.**
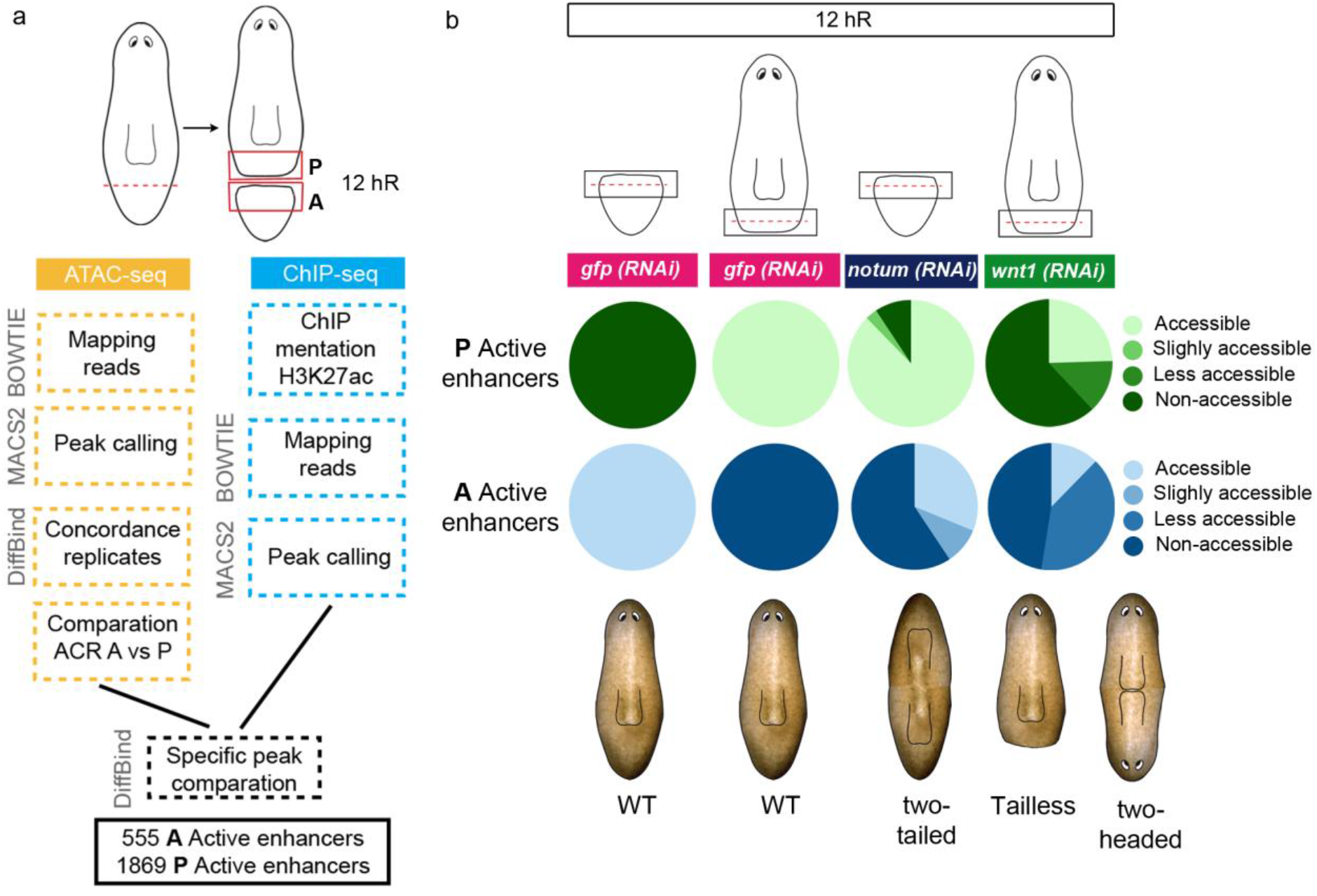
At 12 hours of regeneration anterior and posterior wounds show a specific remodelling of the chromatin. **a** Workflow to identify specific anterior and posterior specific enhancers at 12 hR. Next to each workflow step, the program used is indicated. **b** Accessibility changes of the anterior and posterior specific active enhancers after *notum* and *wnt1* inhibition at 12 hR are represented in percentages in pie charts. Schematic illustration shows the representative phenotypes observed during regeneration after each gene inhibition. hR, hour of regeneration.

Silencing of *notum* or *wnt1* during planarian regeneration produces a shift in polarity, giving rise to anterior tails in *notum* (RNAi) animals (20,42) and posterior heads after *wnt1* (RNAi) (21,22,43). With the aim to analyse the chromatin changes occurred during anterior and posterior specification, we performed ATAC-seq of *notum* (RNAi) anterior wounds and *wnt1* (RNAi) posterior wounds, both at 12 hR. We analysed the state of the anterior and posterior putative active enhancers previously found to be specifically open in anterior or in posterior in those RNAi samples. The result shows that in *notum* (RNAi) anterior wounds only 12.3 % of the anterior putative active enhancers were open, while the rest of them were closed or reduced their accessibility (Figure 1B and Table S1, https://compgen.bio.ub.edu/PlanNET/). Moreover, 87.7 % of the posterior putative active enhancers were now accessible in *notum* (RNAi) anterior wounds. In *wnt1* (RNAi) posterior wounds only 24.5 % of the posterior putative active enhancers were open and the rest were closed or had decreased its accessibility. Furthermore, 31.4 % of the anterior putative active enhancers appeared now open in *wnt1* (RNAi) posterior wounds and 9.5 % became more accessible (Figure 1B and Table S1). The finding that in *notum* and *wnt1* (RNAi) wounds the accessibility of the chromatin changes as soon as 12 hR demonstrates that few hours after amputation, much before the first anterior or posterior markers appear (around 48hR) (44), the chromatin structure of the cells in the wound has already changed according to the polarity of the pre-existing tissue.

Furthermore, the results show that inhibition of the key elements of the anterior and posterior organizers, *notum* and *wnt1*, respectively, produces a very early change of the chromatin structure, suggesting that both elements trigger the specific anterior or posterior program through the regulation of chromatin remodelers. The chromatin changes in *notum* (RNAi) wounds with respect to wt, are much stronger that in *wnt1* (RNAi). This could reflect an earlier role of *notum* in specifying polarity. But it could be also explained because in our experimental conditions *notum* (RNAi) animals polarity is changed, they became two-tailed, while most of *wnt1* (RNAi) animals will became tailless, showing no posterior regeneration, which is the mild *wnt1* (RNAi) phenotype (21,43), and only 10% change polarity and become two-headed.

### Homeobox TFs motives are found enriched in Cis Regulatory Elements of genes downregulated in *wnt1* RNAi planarians and, in turn, contain TCF binding sites

To identify CREs that could be regulated by the cWNT pathway during posterior regeneration we first performed an RNA-seq of controls and *wnt1* (RNAi) posterior wounds (0 to 72 hR) to find the genes downregulated after cWNT pathway inhibition. We performed differential expression analysis (padj<0,05, fc>0,5) at each time point (Table S2). 2129 genes were found to be differentially expressed at any time point; among them 712 genes were down-regulated in *wnt1* (RNAi) planarians with respect to controls (Figure 2B and Table S2). The posterior Homeobox genes are found in the list of down-regulated genes (*Smed-hox4b, Smed-post-2c Smed-post-2b, Smed-lox5a* and *Smed-lox5b*), as well as the posterior Wnts11 (*wnt11-1* and *wnt11-2*) (43, 45), the posterior Frizzled *fzd4* and *sp5*, the TF recently found to mediate the evolutionary conserved role of cWNT in axial specification (46). Interestingly, the TFs required for regeneration of longitudinal and circular fibers, *myoD* and *nkx-1*, respectively (19), are also found among the *wnt1* RNAi downregulated genes, which could account for the rounded shape of the Tailless posterior tip.

**Figure 2.**
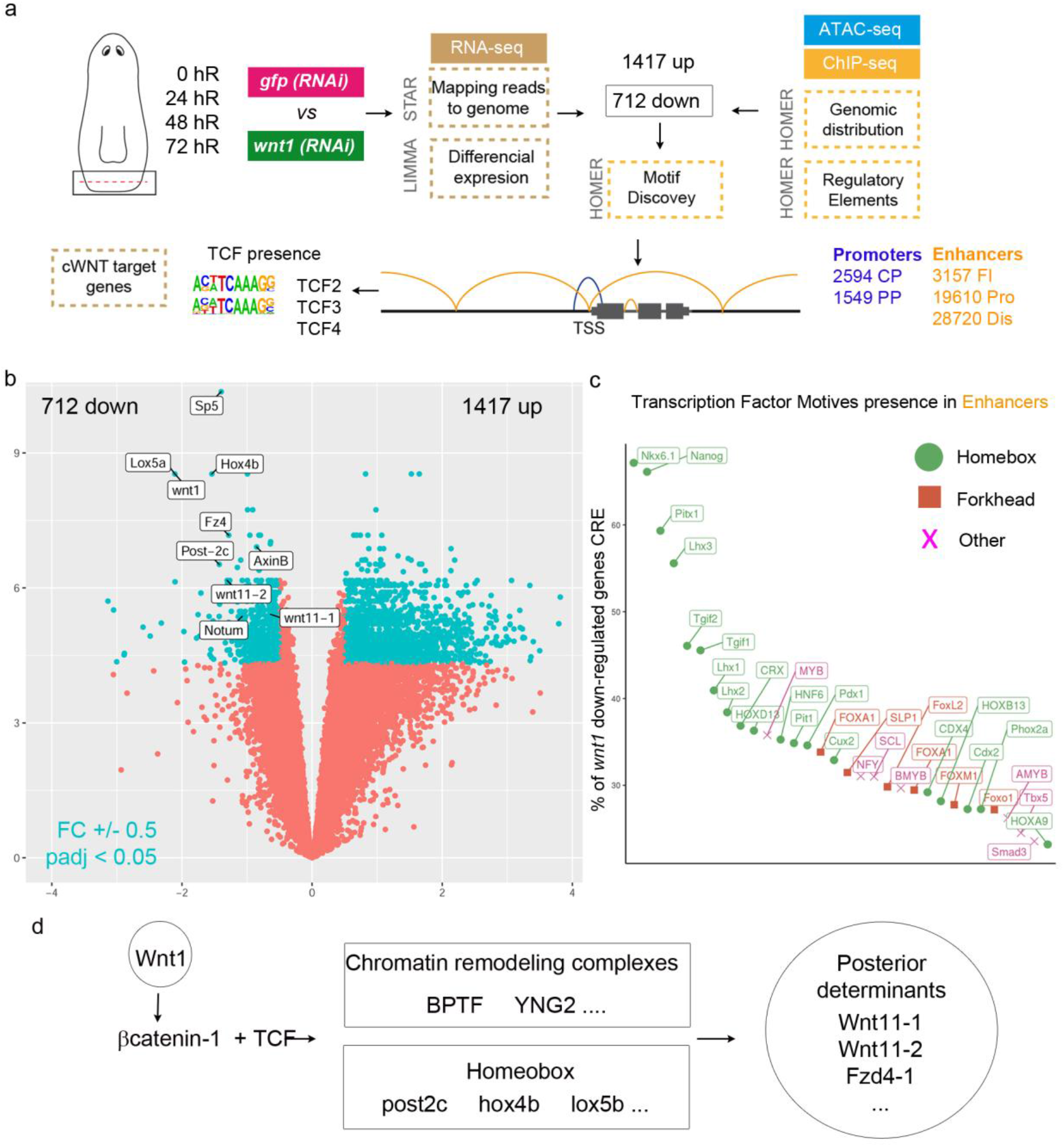
Cis-regulatory regions found in genes down-regulated after *wnt1* RNAi. **a** Workflow to identify differentially expressed genes, cis regulatory elements (CRE) and transcription factors associated to *wnt1* (RNAi). Next to each workflow step, the program used was indicated. Motif discovery for TCF binding site were specifically performed in down-regulated *wnt1* (RNAi) genes. **b** Volcano plot shows the down- and up-regulated genes after *wnt1* inhibition which present fold change (FC) +/- 0.5 and pValue adjusted (padj) < 0.05. **c** Motif enrichment of the CRE of *wnt1* (RNAi) down-regulated genes, showing an enrichment for homeobox motives. **d** Schematic illustration of the proposed genetic program activated by *wnt1* in posterior wounds.

To analyse the CREs of the *wnt1* RNAi down regulated genes we first identified the CREs found in planarian wounds (0-48hR) using the previous ATAC-seq and CHIPmentation samples, in addition to new ATAC-seq analysis of 0 and 48 hR wounds (https://compgen.bio.ub.edu/PlanNET/planexp). We classified the CREs found in Promoters or Enhancers according to their position with respect the Transcriptional start site (TSS) (Figure 2A) (defined in Material and Methods section). Promoters were classified in Core Promoters (CP) and Proximal Promoters (PP), and we identified 2594 and 1549 of each, respectively. Enhancers were classified in First Intron (FI) Proximal (Pro) and Distal (Dis), and we identified 3157, 19610 and 28720 of each, respectively.

Using HOMER we could analyze the presence of TF binding motifs in the CREs of the 712 genes down-regulated in *wnt1* (RNAi) wounds. The result shows that the overrepresented motifs mainly bind Homeobox TFs (Figure 2C). Considering that posterior identity is specified by the cWNT signalling (Wnt1-βcatenin-1–TCF) (47), we then searched for the CRE containing a TCF binding site. We found 167 genes containing a TCF binding site in the enhancer, 17 of which also showed a TCF motif in the promoter (Table S3). Among them, we found the genes already known to be involved in P specification: posterior Hox genes (*lox5b, hox4b, post2c*) (26), *sp5* (46), *axinB* (24) and *tsh* (37,38), indicating that they are direct targets of the Wnt1-βcatenin-1–TCF signalization. We also combined our RNA-seq data with the RNA-seq of *βcatenin-1* RNAi animals already reported (46). This strategy ended up with 42 genes (Table S2), which included most of the posterior genes already found to possess a TCF binding site, further supporting the direct role of these candidates in specifying posterior through the cWNT signalling. A new web-tool has been developed in order to search for transcription factor binding sites and to explore the newly predicted regulatory elements (https://compgen.bio.ub.edu/PlanNET/tf_tools).

As exposed, several genes downregulated in *wnt1* (RNAi) wounds showed a TCF binding site. However, the overrepresented motives found in *wnt1* (RNAi) downregulated genes are not TCFs but Homeobox. These results suggest that the Wnt1-βcatenin-1-TCF signal could directly activate the expression of Homeobox TFs, which in turn will activate the determinants of the posterior fate. In effect, analyzing the list of genes with TCF binding site we found several Homeobox TFs downregulated after *wnt1* (RNAi) in addition to the ones already reported (hox4b, lox5b or Post2c), as BARHL2 or NKX6-2. Among the *wnt1* (RNAi) downregulated genes containing TCF motives we also found chromatin remodelling complexes, as BPTF, a nucleosome-remodeling factor (48), and YNG2, which acetylates nucleosomal histone H4 and H2A (49)(Figure 2D). This result agrees with the rapid changes in chromatin conformation that we have observed at 12h of regeneration.

Overall, we have identified new genes, TFs and CREs participating in the specification of posterior identity; some of them were already known to specify posterior, validating our strategy, and many of them are new elements of the Wnt1 gene regulatory network. Our data suggests that Wnt1-βcatenin-1-TCF signal directly activates chromatin remodelling complexes and Homeobox genes which in turn regulate the determinants of posterior specification.

### FOXG could regulate *wnt1* transcription and specify posterior identity through binding to a Cis-regulatory element (CRE) found in *wnt1* first intron

Taking advantage of the previous analysis that allowed the mapping of CRE in *S. mediterranea* genome (39), we sought to investigate the presence of CRE in the *wnt1* locus, in order to understand the regulation of its expression. We found different evidences that the first intron of *wnt1* presented two enhancers, which were named enhancer 1 (E1) and enhancer 2 (E2) according to their distance to the TSS (Figure 3a and Supp data1). We found that both regions show: (i) a nucleosome free region (ATAC-seq peak) and (ii) that this region is surrounded by histone modifications related with enhancer activity (H3K27ac ChIPmentation) (Figure 3a). Both putative enhancers were located less than 3 kb away from the *wnt1* promoter suggesting that they could regulate its expression (50). Trough motif discovery we analysed the presence of TF binding sites in both regions, and we observed the presence of FOXG binding sites (Figure 3a and Supp data1).

**Figure 3.**
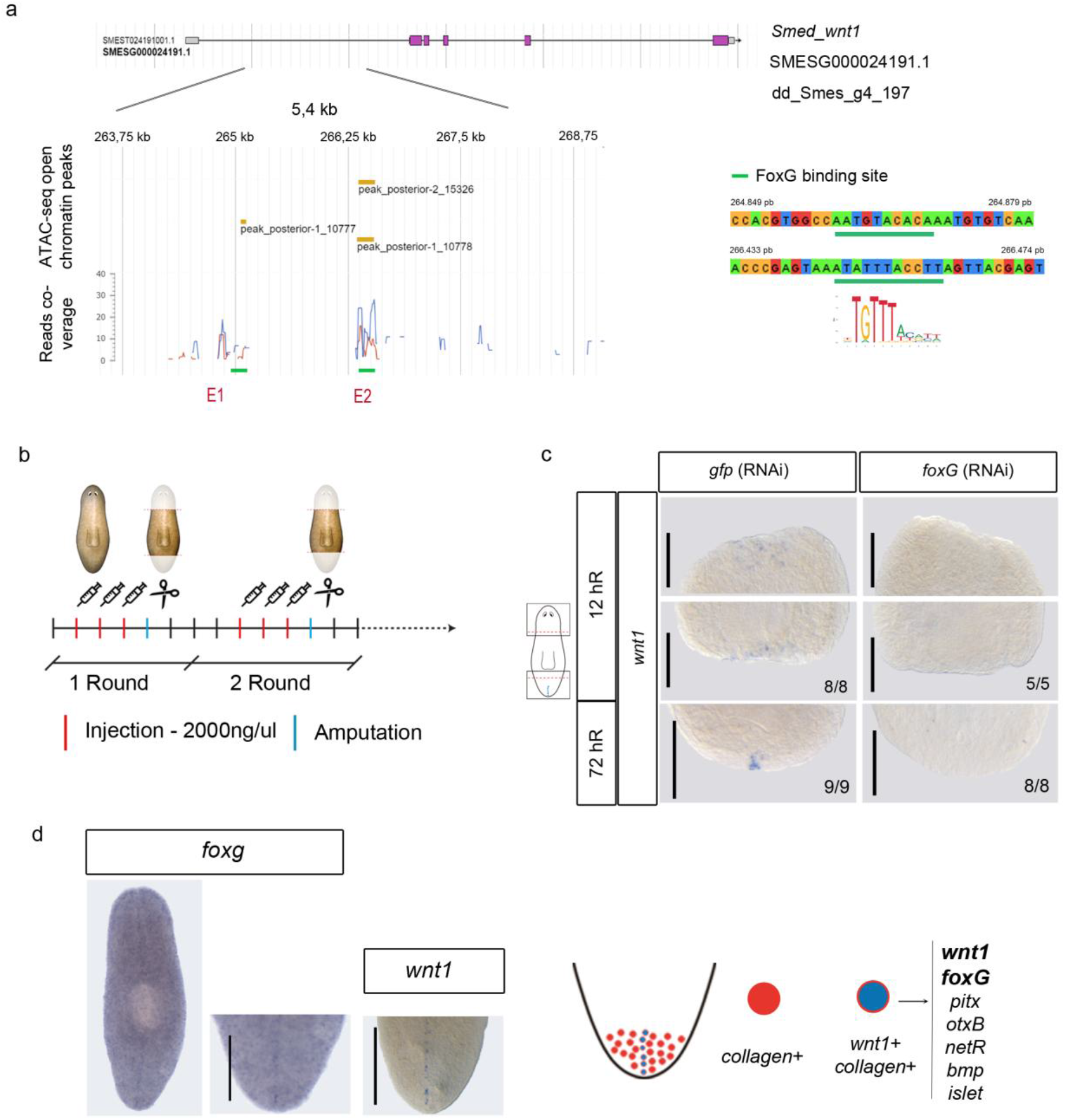
FoxG could bind to a *wnt1* enhancer to regulate *wnt1* transcription. a Schematic illustration of *Smed-wnt1* gene locus, indicating exons (violet boxes) linked by introns (lines). Enhancers are named E1 and E2, FOXG motif (slp1) was present in both enhancers (green line). The ATAC-seq peaks corresponding to the E1 and the E2 are indicated. **b** Schematic illustration indicating the *foxG* RNAi procedure. **c** ISH of *wnt1* in *foxG* (RNAi) animals demonstrate its absence in regenerating blastemas both at 12 and 72hR. Schematic illustration of wnt1 in intact animals and the analysed zones (squares) was added. **d** WISH of foxG in intact animals shows its expression in the posterior midline, as *wnt1*. Single cell analysis performed by (41) showed six genes (top 16%) over represented in posterior organizing *wnt1+* cells. Among them *foxG* was found. Scale bar: 100 μm in c; 200 μm in d.

To further investigate whether FOXG could be regulating *wnt1* expression we inhibited it by RNAi in regenerating animals (Figure 3b). ISH of *wnt1* in *foxG* (RNAi) animals demonstrated that it was absent both at 12hR and 3dR posterior wounds, indicating that *foxG* is required for both the early (stem cell independent) and the late (stem cell dependent) phase of *wnt1* expression (Figure 3c). *foxG* was also necessary for the expression of *wnt1* in the anterior 12hR wounds (Figure 3c). This result is important, since it is the first gene reported to date that regulates the early *wnt1* stem cell independent expression that occurs few hours after amputation in any wound. Furthermore, inhibition of *foxG* in intact animals also lead to the disappearance of *wnt1* expression (Figure S1), supporting its general role for the expression of *wnt1* in planarians, possibly by binding to the enhancer found in the first intron of *wnt1*.

In agreement with a direct role of *foxG* in regulating *wnt1* expression and posterior specification, ISH analysis shows that *foxG* is expressed in the posterior dorsal midline, as described with *wnt1* expression (Figure 3d). *foxG* is also expressed in cells along the D/V margin, and in some scattered cells in the dorsal and ventral part of the animals. Single Cell Sequencing (SC-seq) databases analysis indicates that those cells could be muscular and neuronal (Figure 3d and Figure S21). To note, *foxG* is one of the specific genes found in muscular *wnt1+* cells of the posterior midline (*wnt1*+ and *collagen*+) in intact animals and in posterior regenerating blastemas at 72 hR (51,52), further supporting its essential role in the specification of the posterior organizer (Figure 3d).

If *foxG* is required for *wnt1* expression, then regenerating *foxG* (RNAi) animals should show a phenotype related with the malfunction of the posterior organizer. Accordingly, we found that 70% of *foxG* (RNAi) animals presented a rounded posterior blastema, resembling the tailless phenotype obtained in the mild *wnt1* RNAi phenotype (Figure 4a). Analysis of the central nervous system and the digestive system by anti-arrestin (3C11) and anti-βcat-2 immunohistochemistry, respectively, demonstrated that these 70% animals are tailless (Figure 4b and Figure S2b). They show a fusion of the posterior nerve cords and intestine branches in U shape, as it has been described after inhibition of other key posterior genes such as *wnt1* (21,43), *wnt11-2* (29,43), *islet* (35,36) or *pitx* (36). Furthermore, ISH with posterior markers, which we demonstrate in the previous section are cWNT target genes (*fz4*, *post2d*, *sp5* and *hox4b*) demonstrates that they are downregulated in posterior *foxG* (RNAi) blastemas at 3 dR (Figure 4b). Eventually some animals showed the strongest two-headed phenotype (Figure 4a). Synapsin analysis demonstrates the appearance of a posterior brain in these animals (Figure 4c).

**Figure 4.**
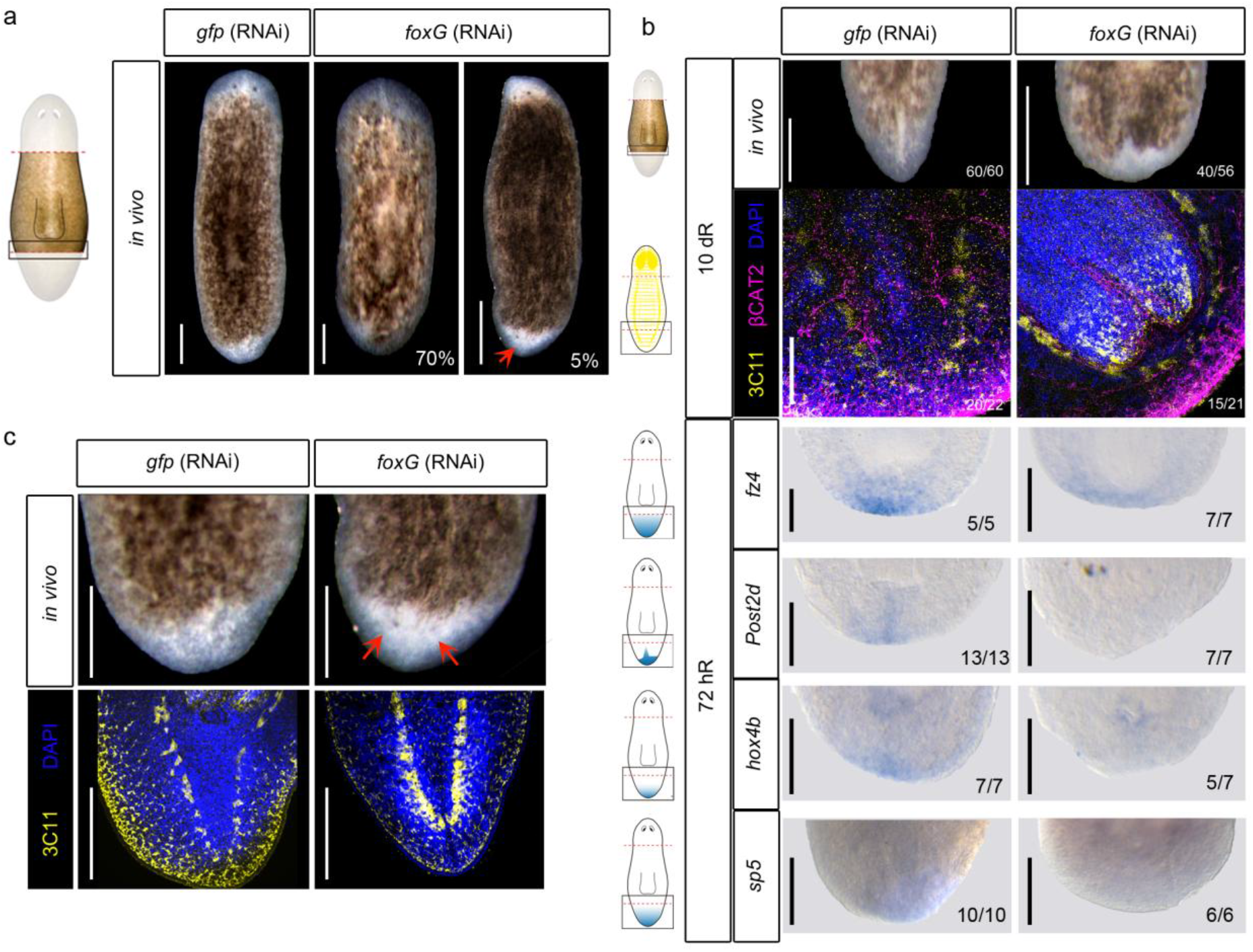
*foxG* RNAi phenocopies *wnt1* inhibition. **a** *in vivo* phenotypes after *foxG* (RNAi). **b** Immunostaining using α-SYNAPSIN (3C11) (neural system) and α-βCAT2 (digestive system) reveal rounded ventral nerve chords in *foxG* (RNAi) compared to peak shaped *gfp* (RNAi) animals. Nuclei are stained in DAPI. WISH of posterior markers in regenerating *foxG* (RNAi) animals demonstrated a reduced expression. Schematic illustrations of posterior markers were added. **c** Immunostaining using α-SYNAPSIN (3C11) (neural system) reveal a posterior brain in the *foxG* (RNAi) Two-headed animals. Nuclei are stained in DAPI. Posterior eyes are indicated with a red arrow in a and c. Posterior eyes are indicated with a red arrow in a and c. Scale bar: 100 μm in a, immunostaining in b and c: 200 μm in WISH in b.

All together, these results demonstrate that *foxG* is a new element of the posterior organizer. Our data indicates that *foxG* is upstream of *wnt1* because inhibition of *foxG* suppresses *wnt1* expression in all stages and tissue regions, and because *wnt1* (RNAi) animals do not show a decrease in *foxG* expression (Table S2 and Figure S2c).

Although our data does not demonstrate the direct binding of FoxG to the enhancer found in the first intron of *wnt1*, we looked for evidences showing the evolutionary conservation of this enhancer. Interestingly, we found that the position of *Schmidtea mediterranea* intron 1 is highly conserved through evolution (Fig 5). *wnt1* genes present a variable number of introns, however, the first intron, which contains the FoxG binding site, maintains a conserved position in all the genomes analysed (Fig 5a and Table S4). Furthermore, the analysis of reported ATAC-seq data demonstrates the existence of chromatin open regions in this intron (Fig 5 and Table S4). More importantly, in *Drosophila melanogaster* there is a CHIPmentation analysis using the FoxG antibody that demonstrates the binding of DmFoxG (slp1) in the first intron of *Dm-wnt1* (53,54).

**Figure 5.**
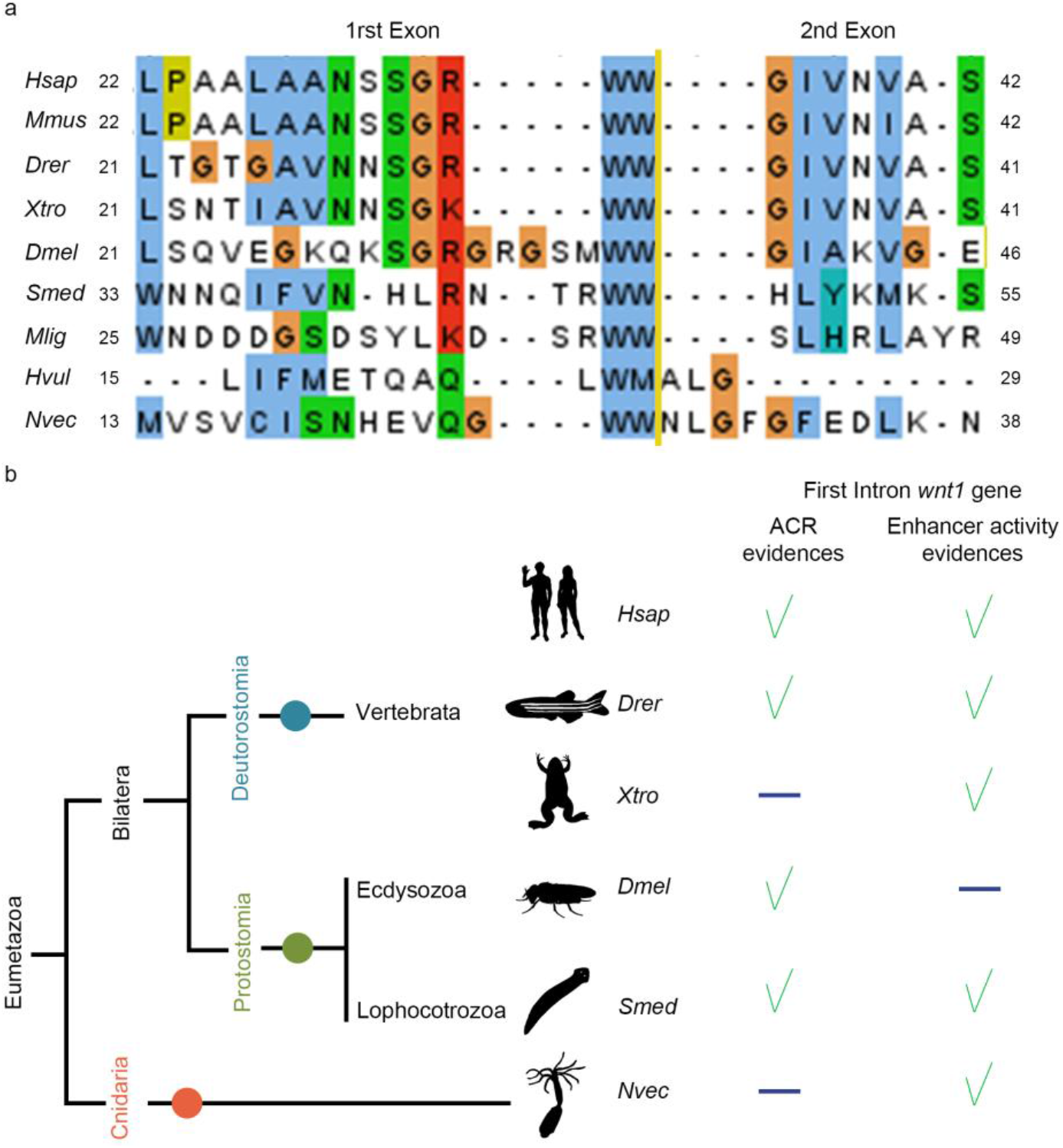
Enhancers located in the first intron of *wnt1* are conserved thought evolution. **a** Alignment of WNT1 amino acid sequences from *Homo sapiens* (*Hsap*), *Mus musculus* (*Mmus*), *Danio rerio* (*Drer*), *Xenopus tropicalis* (*Xtro*), *Drosophila melanogaster* (*Dmel*), *Schmidtea mediterranea* (*Smed*), *Macrostumum ligano* (*Mlig*), *Hydra vulgaris* (*Hvul*) and *Nematostella vectensis* (*Nvec*) showing high level of conservation on the position of intron 1. Yellow line shows the separation between the first and the second exon. **b** Schematic summary of accessible chromatin regions (ACR) and enhancer activity evidences in the first intron of *wnt1* genes in different metazoan species. Green tick indicates evidences and blue line indicates no available data.

Thus, our data supports the hypothesis that in planarians FoxG could directly regulate *wnt1* expression, and the early establishment of the posterior organizer, through binding to a first intron enhancer.

## Discussion

### Dynamics of genomic and transcriptomic changes occurring during posterior identity specification

The plasticity of planarians is providing insightful data about the mechanism underlying regeneration. Several studies now demonstrate that the anterior and the posterior tips of planarians function as organizers, a term that has been traditionally used in the field of embryonic development. The finding of adult organizers in other regenerating animals as hydra, zebrafish or xenopus tadpoles supports the idea that the formation of organizers could be a general mechanism that confers regenerative properties (55–57) There are common features in the reported examples: 1) the cells that function as organizers are non-proliferative and are located in the periphery of the early blastema, 2) the organizing activity relies of the cWNT signal (55,56,58). These properties are also accomplished by planarian organizers. A difference between planarians and other bilaterian models of regeneration is that planarians can completely regenerate a new axis from both ends, anterior or posterior, independently of the fragment amputated. This plasticity, together with the use of genomic and transcriptomic high throughput techniques, has allowed us to compare the genomic changes occurring during anterior or posterior specification in the same field of original cells. Our data indicates that the establishment of the appropriate identity in a planarian wound could follow the following 3 steps (3-step model in Figure 6). 1^st^) remodelling of the chromatin, which must occur very early after a cut, even before the appearance of any anterior or posterior marker. We demonstrate that the regulation of the cWNT signal is fundamental for this remodelling. We have seen a strongest contribution of *notum* in anterior wounds than *wnt1* in posterior wounds in this remodelling. However, we cannot conclude that *notum* has a more determinant or earliest effect than *wnt1*, since we only have analyzed one time point, 12 hours after the cut, and furthermore our *wnt1* (RNAi) animals have a milder phenotype (Tailless) than *notum* RNAi animals (Two-tailed). 2^nd^) Remodelling of the chromatin could allow the expression of polarity genes, as the Hox genes, which expression has shown to be dependent on extensive chromatin remodelling in other models (59). 3^rd^) Hox genes among others would allow the transcription of the Posterior determinants, which are the effectors required to differentiate tail structures.

**Figure 6.**
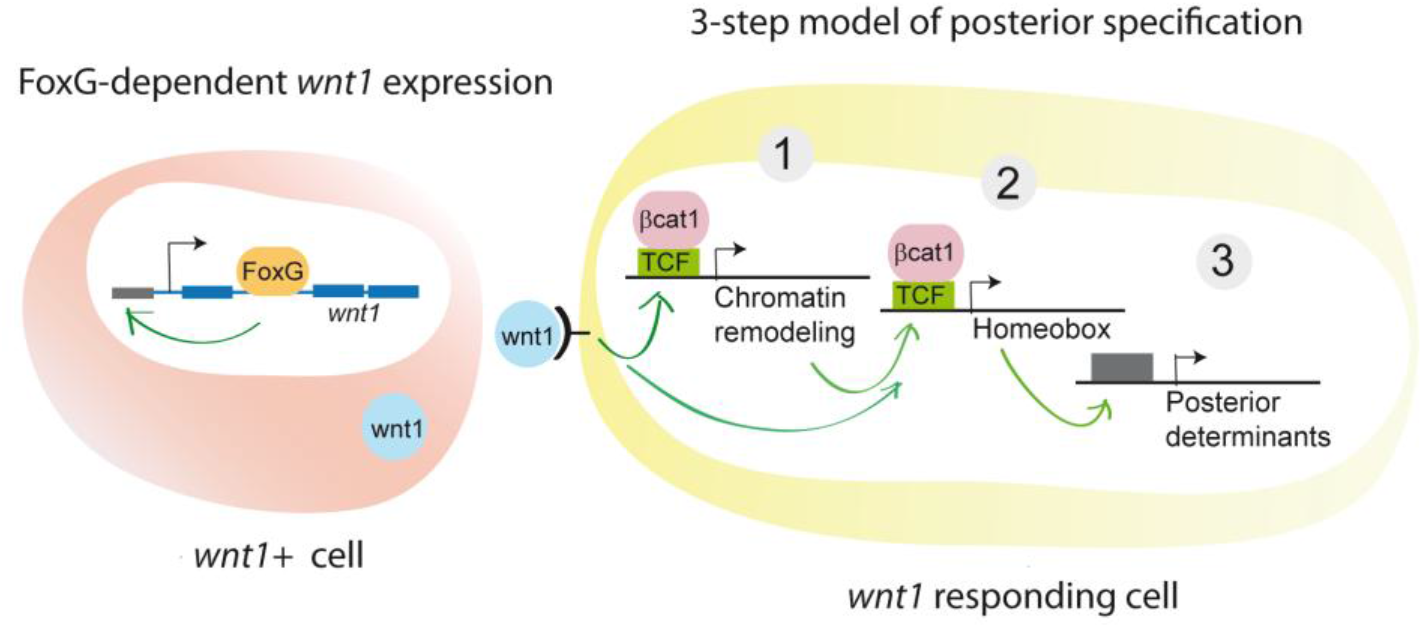
3-step model of posterior specification.

The early change in the genomic landscape found in each regenerating tip, together with the finding of several chromatin remodelling proteins down-regulated in *wnt1* (RNAi) genes showing a TCF binding site (refs) indicates that in planarians the Wnt/β-catenin pathway specifies cell fate through regulating chromatin structure and reprogramming as described in other contexts (60). It is important to note that the early expression of *notum* and *wnt1* is stem cell independent (22), supporting the important role that reprogramming could have at this early stages. Importantly, our data restricts the timing when this chromatin remodelling it is happening, which must be earlier than 12 hours after the cut. Thus, a novelty of the proposed 3-step model is that, in contrast to the results found in previous transcriptomic analysis in planarians, in which injury-specific transcriptional responses emerged 30 hours after injury (44), we have observed that changes occurring in the chromatin of cells in each wound are wound-specific and occur few hours after the cut. These rapid changes at genomic level have been visualized thanks to the genomic analysis restricted to the cells in the wound region and at a very early time point (12 hR), much before the appearance of any polarity signal.

Our RNA-seq analysis agrees with previous transcriptomic studies, since we have found that inhibition of *wnt1* leads to the deregulation of a large number of genes at late time-points (48-72h), corresponding to the previously called injury-specific transcriptional response (44). However, the second novelty of our 3-step model is that through the identification of the CREs in *Smed* genome, we could observe only few *wnt1* (RNAi) down-regulated genes containing TCF motives in their CREs, and many of them correspond to chromatin remodelling complexes and homeobox proteins. This result suggests that Chromatin remodelling proteins and Homeobox genes are direct targets of WNT1 that afterwards will activate the transcription of the posterior determinants. The Hox genes *post2c, lox5a/b* and *hox4b* are specifically expressed in posterior, although regenerative deflects have only been see after *lox5a* (RNAi) (45, 46). Not only in planarians but also in other whole body regenerating animals as acoels and hydra Hox genes and *sp5*, which also shows a TCF motive in its CRE, are the conserved set of cWNT targets that mediate the patterning of the primary body axis (46,58).

According to our data, *wnt11-1* and *wnt11-2*, which are required to regenerate a proper tail but whose inhibition never produces a shift in polarity (21,22,43), are down-regulated at late stage in *wnt1* (RNAi) animals, and do not show a TCF binding motive in their CRE, suggesting that they could be part of what we have called the Posterior effectors. Supporting the late role of *wnt11-1* and *wnt11-2*, their silencing inhibits the late *wnt1* expression but not the early one (43). The same situation is found with the posterior expressed WNT receptor, *fzd4*. In this case, it could be that the expression of *fzd4* is mediated by the first *wnt1+* cells forming the posterior organizer, in accordance with the idea that organizers evocate the surrounding tissue; a first organizer action would be to prepare the tissue to make it competent to itself (2).

### Establishment of the posterior organizer requires a FOXG mediated WNT1 signal

The fundamental role of the Notum-Wnt1 antagonism in establishing the identity of a wound has been widely demonstrated through functional and expression analysis (20–22,43). The proposed 3-step model assumes this antagonism and presupposes that the remodelling of the chromatin is different in anterior and posterior wounds because *notum* is expressed at higher levels in anterior and *wnt1* is expressed at higher levels in posterior. Genes required for the late expression of *notum* and *wnt1*, localized in the midline, have been identified. *foxD* and *zicA* RNAi animals do not show the late expression of *notum* and do not regenerate a proper head (30–32); *islet* and *pitx* for RNAi animals do not show the late expression of *wnt1* and are Tailless (35,36,61). However, the regulation of the expression of *notum* and *wnt1* in a salt and pepper manner in early wounds and their final restriction to the anterior and posterior pole, respectively, remained unsolved.

Thanks to the annotation of the CREs in the planarian genome, we could identify an enhancer in the first intron of *wnt1* gene which showed FOXG binding motives. We propose that these CREs are general enhancers required for *wnt1* expression in planarians. First, they are localized in the first intron, which is a region frequently enriched in regulatory elements (62–65); second, we have found that *foxG* is necessary for *wnt1* expression in any context. *foxG* inhibition suppresses the early and the late phase of *wnt1* expression during regeneration, as well as its expression during homeostasis. And third, it could be evolutionary conserved, which further supports its relevance. We have found that the position of intron 1 in all *wnt1* genes studied from different metazoans species is extremely conserved. Furthermore, there are several genomic studies that demonstrate the existence of open regions in this intron, and a Chip-sequencing analysis with the FoxG antibody in *Drosophila* demonstrates that foxG binds to *Dm-wnt1* (wingless) intron 1. We hypothesize that the binding of FoxG to the intron1 of *wnt1* to regulate its expression is ancestral and has been conserved through evolution. Genomic studies in different regenerative species have identified different sets of TFs as regulators of cWNT genes during regeneration. In Drosophila, injured imaginal disc required ‘regenerative enhancers’ to trigger *wingless* expression and the regeneration process (66–68). During *Hydra* head regeneration, an enhancer collection become accessible allowing the expression of cWNT genes in head organizing cells (69,70). In acoels, *egr* is expressed after amputation triggering the expression of *wnt3*, which participates in the posterior specification (67, 71). As recently proposed, it could be that enhancers may be maintained as part of conserved gene regulatory network modules over evolution (72). In this respect, further studies are required to analyse the evolutionary conservation of the enhancers found in the first intron of *wnt1*.

RNAi inhibition of *foxG* suppresses both early and late *wnt1* expression after amputation, and, consequently, animals became Tailless. Importantly, in a low percentage, *foxG* (RNAi) animals regenerated as Two-headed. A shift in polarity is a phenotype only found after *βcatenin1* or *wnt1* RNAi (21,24,28), but never after *islet* or *pitx* RNAi. Thus, the finding of Two-headed *foxG* (RNAi) animals, suggests that inhibition of the late phase of *wnt1* prevents the regeneration of a tail, but that inhibition of the early phase is required to shift polarity. This idea is supported by the reports on the role of Hh signal in planarians. Activation of the hh signal is also required for the early phase of *wnt1* expression and in a low percentage animals become two-headed (73,74). According to these data, Hh could mediate its early role in polarity establishment by regulating *wnt1* expression through foxG activation, as it has been reported in zebrafish and mouse, where Hedgehog signalling contributes to Foxg1 induction and integration of telencephalic signalling centres (75,76).

The planarian posterior organizer is defined by the expression of *wnt1* in differentiated muscular cells. However, the cells in the organizer must integrate a signalling network which includes several genes which are not cell specific, and simultaneously they must be integrated in a diverse a dynamic cellular context. Our study has provided some light to this genetic and cellular context of the planarian posterior organizer. We have found *foxG* as a gene essential for *wnt1* expression, which inhibition phenocopies *wnt1* RNAi. However, *foxG* is not specifically expressed in *wnt1+* muscular cells, but it is also expressed in neurons and in progenitor cells. In fact, we have found that the binding site for FoxG (SLP1) was also notably enriched in enhancers of *wnt1* (RNAi) down-regulated genes. Thus, FoxG could regulate not only *wnt1* expression in muscular cells but additional *wnt1* regulated posterior genes in other cell types, as neurons. Further studies are necessary to analyze whether FoxG binds to E1 and/ or E2 of *Smed wnt1*, as well as the existence of specific co-factors that confer it a cell-dependent activity.

### Conclusion

The existence of ‘regenerative enhancers’, groups of enhancers that become accessible during regeneration, has been demonstrated in regenerating species as zebrafish and *Drosophila* imaginal discs (66,77–80). The plasticity of planarians, which make possible the comparative study of anterior and posterior regenerating wounds originating from the same field of cells, has allowed the identification of ‘regenerative enhancers’ specifically associated to the posterior specification. The data presented in this study suggests that the formation of the posterior organizer could be working as a chain reaction. A first differential signal in the wound according to the polarity of the pre-existent tissue (which could be related to Hh or other neural signals) (73,74) would lead to the rapid resolution of the Notum-Wnt1 antagonism, which in posterior wounds will maintain Wnt1 and suppress Notum. At this point, which must occur during the first 6 hours the program to become posterior has already started, setting up chromatin changes specific to the posterior pole and dependent on cWNT activation. Chromatin conformation changes would allow the subsequent expression of a specific set of TFs that will turn on the tail effectors.

Organizers or organizing centers are required for growth and pattern of a new structure and are well studied during embryonic development. However, in whole-body regenerating animals as planarians or hydra, organizers are also found in adults. Muscular *wnt1+* cells are found in the midline of posterior wounds and in the tail of planarians during homeostasis (22,29), and we have found that in both contexts *wnt1* expression depends on *foxG*. However, homeostatic and regenerating *wnt1+* cells have different properties, since inhibition of *wnt1* or *foxG* during homeostasis never produces a shift in polarity. After an amputation, when new tissue must be regenerated, there must be a time window when everything is possible. According to the signal received by the cells in the wound the identity of the organizer will be decided. Importantly, not only the identity but the presence of an organizer, which means the possibility to regenerate or not, will be determined. This was shown in Liu et al. when modulating the cWNT provided regenerative capacity to planarian species that are not able to regenerate a head in nature (81). These results indicate that the ability to form an organizer is linked to the ability to regenerate. Thus, understanding the formation and function of Organizers is key to understand adult regeneration.

Finally, our results demonstrate the power of genome wide approaches to further understand the genetics of regeneration. With the aim to share the results obtained in this study and to facilitate their further analysis to the scientific community, we have created a new open platform to query and interpret all transcriptomic and genomic results obtained (https://compgen.bio.ub.edu/PlanNET/planexp). Furthermore, an additional web-tool has been developed in order to search for transcription factor binding sites and to explore the newly predicted regulatory elements (https://compgen.bio.ub.edu/PlanNET/tf_tools).

## Material & Methods

### Planarian husbandry

*Schmidtea mediterranea* clonal strain BCN-10 animals were starved for at least 7 days prior any conducted experiment. Asexual animals were cultured in glass containers and Petri dished for experiments in PAM water (82) at 20°C the dark. Animals were regularly feed twice per week with organic cow liver (83).

### RNAi experiment design

For RNAi, double strand RNA (dsRNA) was synthesised by *in vitro* transcription (Roche) using PCR-generated templates with T7 and SP6 flanking promoters. Precipitation step was carried using ethanol, followed by annealing and resuspension in water. (92). dsRNA (3 × 32.2 nl) was injected into the digestive system of each animal on 3 consecutive days (1 round). For *wnt1* RNA-seq samples, inhibited and control animals were injected one round at 1500 ng/μl and amputated at post-pharyngeal level. Then, studied pieces were soaked in dsRNA diluted in PAM water for 3 hours in the dark. For *wnt1* and *notum* ATAC-seq samples, inhibited and control animals were injected two rounds at 1000 ng/μl and amputated at pre- and post-pharyngeal level. *foxG* RNAi regenerating animals were inhibited two rounds and amputated at the end of each round; and intact animals were inhibited for three consecutive rounds. All control animals were injected and/or soaked with dsRNA of GFP.

### Assay for transposase-accessible chromatin sequencing (ATAC-seq)

ATAC-seq samples were obtained from the wound region of wild type, *notum* (RNAi), *wnt1* (RNAi) or *gfp* (RNAi) samples. Planarian mucous was removed by washing in 2%L-Cystein (pH7) for 2’. Afterwards, animals were transferred in a petri dish with CMFH (2.56mM NaH2PO4×2H2O, 14.28mM NaCl, 10.21mM KCl, 9.42mM NaHCO3, 1%BSA, 0.5%Glucose, 15mM HEPES pH 7.3). Planarians were placed in Peltier Cells at 8°C to amputate the wound region (the blastema and post blastema region posterior to the mouth). Then, transferred to an 1.5 ml Eppendorf tube to be dissociated using a solution of liberase/CMFH (1:10) at RT for 10 minutes. Twenty animals were used per biological replicate. ATAC-sequencing was carried out as first described in (84) and then adapted by (85).

### ChIPmentation

ChIPmentation combines ChIP with library preparation using Tn5 transposase, similar to ATAC-sequencing. ChIPmentation samples were obtained from the wound region of wild type animals. Planarians were placed in Peltier Cells at 8°C to amputate the wound region (the blastema and post blastema region posterior to the mouth). Then, wounds were transferred to a Petri dish containing 1M MgCl2 solution, for 15-30’’ rocking at RT. PBS 1X was added to remove salts. Blastemas were fixed with formaldehyde 1,85% for 15’ rocking, at RT. Glycine was added to obtain a final concentration of 0.125M to quench formaldehyde, for 5’ at RT, rocking. Then, blastemas were washed 3X with cold PBS1X. Finally, PBS excess was removed, and samples were stored at −80°C. 2000 anterior and posterior blastemas were used. Groups of 100 blastemas were done at a time. ChIPmentation was carried out as described in (86).

### ATAC-seq and CHIPmentation analysis

Reads were aligned using bowtie1 using −m 3 −k 1 arguments. Bam reads were filtered using a <=100bp insert size threshold to identify nucleosome free regions (NFR) (87). Bam files were converted to bed and then the coordinates were shifted +4 and −5 positions to overcome the Tn5 cut position. MACS2 were used for peak calling and HOMER for motif discovery. Differential binding analysis was carried out using DiffBind (R function).

#### Cis-regulatory elements annotations

Putative cis-regulatory elements (CRE) were annotated over the *Schmidtea mediterranea* genome version S2F2. For this purpose, both ChIP-seq and ATAC-seq data from all collected samples were used. Narrowpeaks over the genome were identified using MACS2 (see ATAC-seq and ChIPmentation analysis section of materials and methods). These peaks were merged using the mergePeaks command of the HOMER software suit. Finally, regions over the genome were classified as either *putative promoters*, or *putative enhancers*, according to their evidences regarding ATAC-seq and ChIP-seq peak coverage. Only those regions called as peaks on at least two samples were considered, and the rest were discarded for the CRE annotation.

Peak regions with only ATAC-seq evidences were classified as *core promoters* (< 100 bp upstream of an annotated TSS) or *proximal promoters* (between 500 and 100 bp upstream of a TSS). Finally, peaks with CHIP-seq evidences were classified as either *proximal enhancers* (within 2000 bp of an annotated TSS) or *distal enhancers* (between 2000 and 10000 bp of a TSS).

#### Motif finding

Putative transcription factor binding sites were identified and annotated on all these enhancer and promoter regions by using the HOMER’s *findMotifsGenome* command, scanning these regions using the motifs provided by the software suite (*known motifs*).

Transcription factor motif overrepresentation analyses were also performed using the *findMotifsGenome* command of HOMER. Motifs were called as overrepresented using an adjusted p-value cutoff of 0.05.

#### Integration with online resources

A new plugin for the PlanNET web service (88), called *tf tools*, was developed in order to integrate the putative CRE dataset with existing planarian resources. A search tool for exploring genes according to the presence or absence of transcription factor binding motifs was developed, and the putative CRE elements were incorporated to the existing gene cards in PlanNET and to our genome browser instance. The website, the source code of the plugin, and downloads for all the annotations, will be released upon publication on https://compgen.bio.ub.edu/PlanNET.

#### RNA sequencing sample preparation and analysis

RNA-sequencing samples were obtained after trough the soaking protocol. At the corresponding time point (0-24-48-72 hours of regeneration), animals were placed in a Petri dish with cold 1%HCl (diluted in water) for 2’ and then transferred to a new Petri dish with cold PBS 1X. Two washes were performed with cold PBS 1X and animals were transferred to cold RNAlater for 20’ placed in ice. Afterwards, planarians were amputated in a Peltier Cell with a clean blade, to obtain the blastemas and post-blastemas. Fragments were washed with RNAlater (Invitrogen) and 50% RNAlater /Trizol. Finally, liquids were removed and 100 μl of TrIizol reagent (Invitrogen) was added. Total mRNA extraction was performed as described in (89). Three biological replicates were used per time point. Each biological replicate was composed by eight animal fragments. Libraries preparation and sequencing was carried out by Centre Nacional d’Anàlisi Genòmic (CNAG).

RNA reads were mapped against the planarian genome version S2F2 (39) using the STAR software tool (90). Lowly expressed genes were filtered by removing genes with less than 1 count-per-million (CPM). Two biological replicates were removed due to ineffective *wnt1* inhibition. Differentially expressed genes were detected using the lima-voom pipeline (91), using an FDR cut-off of 0.05 and a log fold change cut-off of ± 0.5.

#### Whole mount *in situ* hybridization (ISH)

RNA probes were synthesized *in vitro* (Roche) using T7 or SP6 polymerases and DIG- or FITC-modified, purified with ethanol and 7.5M of ammonium acetate, diluted in 25 μl ddH_2_O and adjusted to a final concentration of 250 ng/μl. For colorimetric ISH was performed (92): animals were killed in 5% N-acetyl-L-cysteine (NAC), fixed in 4% formaldehyde (FA), permeabilized with Reduction solution for 5’ at 37°C and stored in methanol at −20°C. Following overnight hybridization, samples were washed twice with 2x SSC with Triton-X (SSCTx), 0.2x SSCTx, 0.02x SSCTx and MABTween. Subsequently, blocking was in 5% Horse Serum (X?) and 0.5% Western Blocking Reagent (Roche) MABTween solution and anti-DIG-POD was used. Antibody was washed for 2 hours followed by NBT/BCIP development.

#### Immunohistochemistry staining

Whole mount immunohistochemistry was performed as (93): animals were killed with cold 2% HCl and fixed with 4% FA at RT. After 4 hours in blocking solution (1% BSA in PBS Triton-X 0.3%), animals were stained overnight at 4°C. Animals were washed extensively with PBSTx, blocked for 2 hours and stained overnight at 4°C. The following antibodies used in these experiments: mouse anti-synapsin (anti-SYNORF1, 1:50; Developmental Studies Hybridoma Bank) and anti-Smed-b-catenin2 (1:1000; (94)). The secondary antibodies used were Alexa 488-conjugated goat anti-mouse (1:400; Molecular Probes; A28175) and Alexa 568-conjugated goat anti-rabbit (1:1000; Molecular Probes; A-11011). Nuclei were stained with DAPI (1:5000).

#### Image acquisition

*in vivo* images were acquired with X. Brightfield colorimetric ISH images were obtained with a ProgRes C3 camera from Jenoptik (Jena, TH, Germany). A Zeiss LSM 880 confocal microscope (Zeiss, Oberkochen, Germany) was used to obtain confocal images of whole-mount immunostainings. Fiji/ImageJ was used to show representative confocal stacks for each experimental condition are shown.

## Data availability

Data set generated is deposit in GenBank with the accession numbers: XXX

## Acknowledgements

We wish to thank all members of the Emili Saló, Teresa Adell and Francesc Cebrià labs for their suggestions and discussion of the results. A thought to the memory of José Luis Gomez Skarmeta, who passed away on September 16, 2020 while this manuscript was in progress; he was an exceptional promoter of scientific collaborations.

## Author Contributions

E.P-C. and T.A. designed the study and wrote the manuscript; E.P-C., M.M-B and S.C-L. performed and analyzed genomic experiments; P.C-C performed foxG RNAi experiments; S.C-L. and M.M-B performed the bioinformatic research; J.F.A supervised the bioinformatic research. M.M-B, M.S.M., G.N.W. and J.L.G-S. performed the Chip-seq experiments and analyzed the results; E.S. and T.A. received the funding and supervised the research.

## Supplementary information

**Figure S1.**
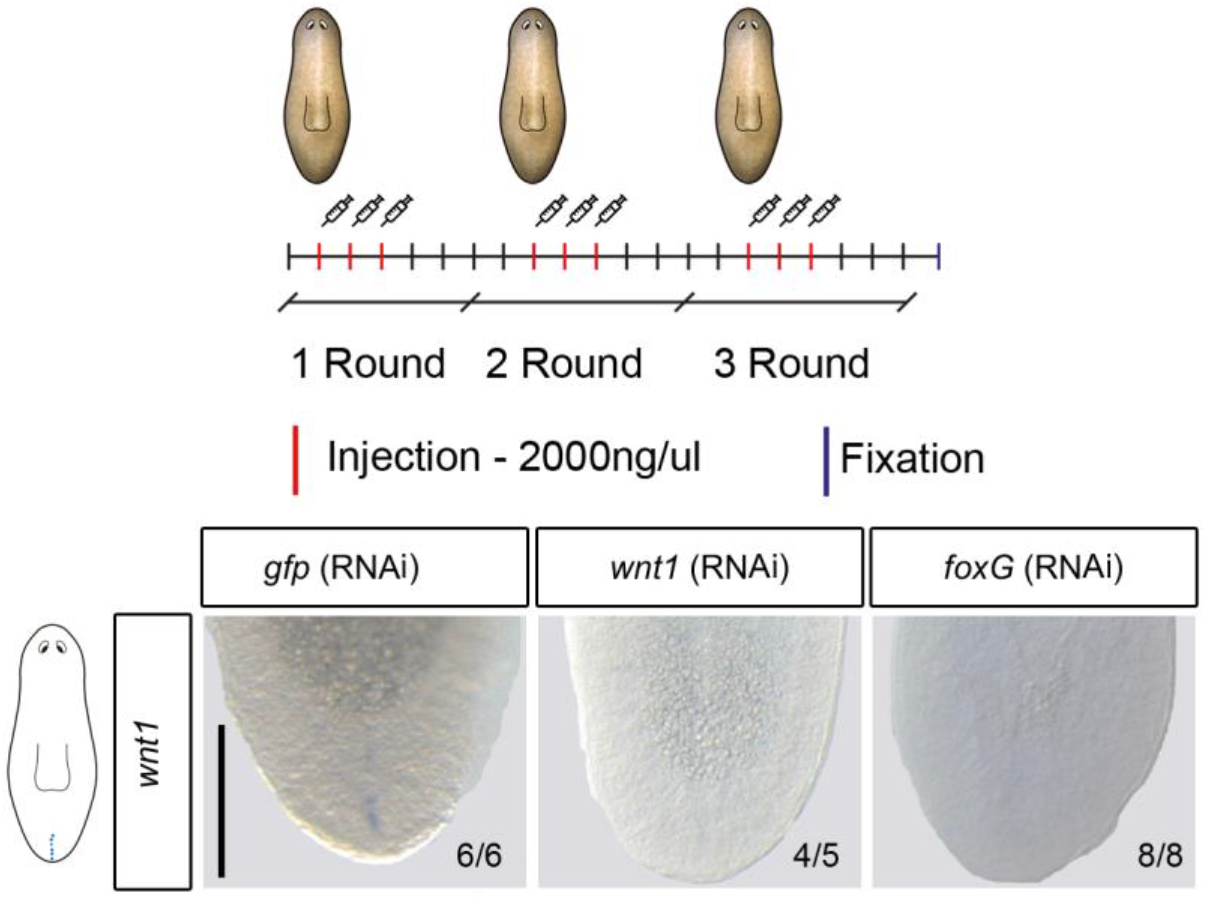
**b** WISH of *wnt1* in intact animals after *wnt1* and *foxG* inhibition reveal its absence. Schematic RNAi design illustration was added.

**Figure S2.**
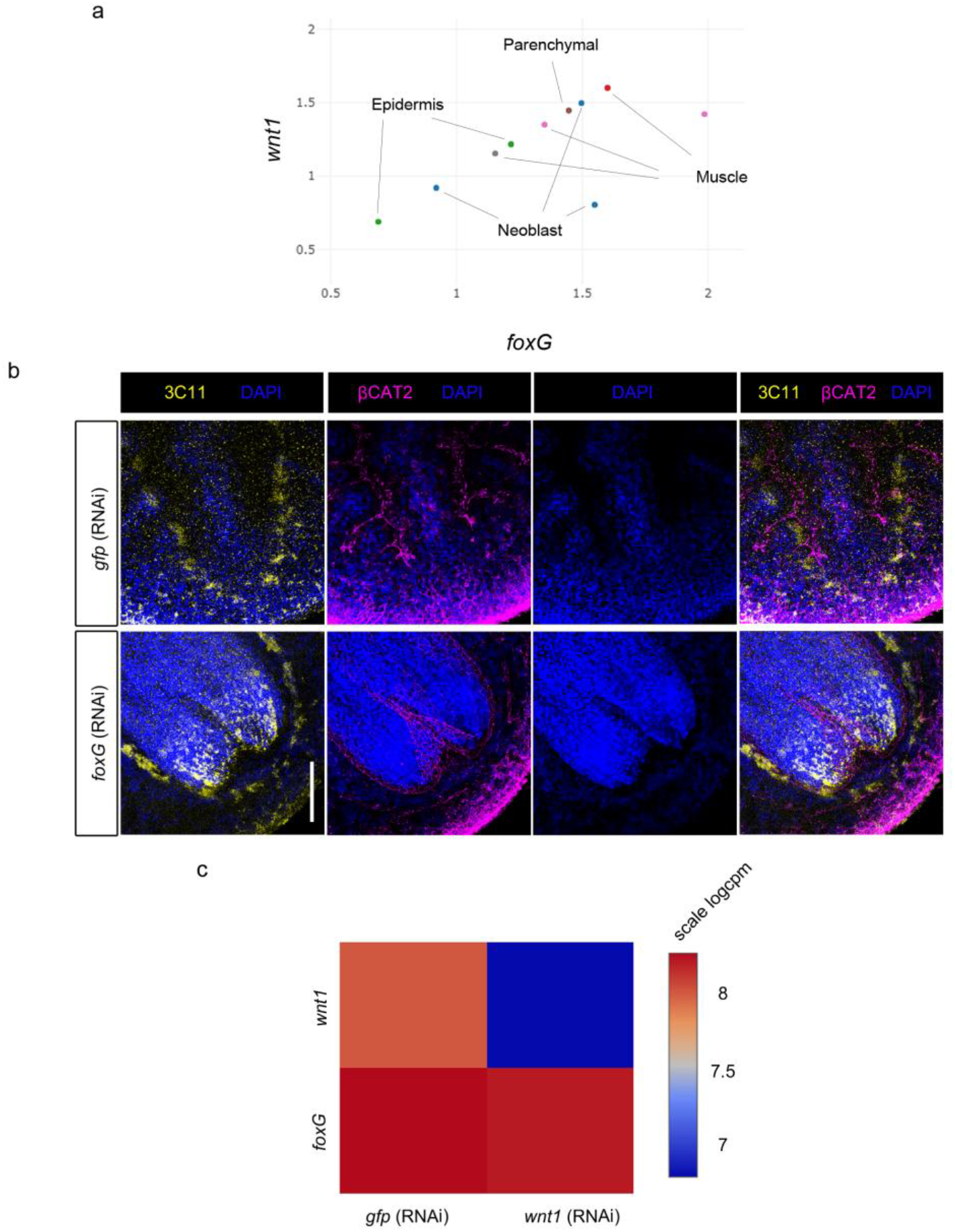
*foxG* (RNAi) animals showed lack of posterior regeneration. **a** SC-seq data from (95) demonstrates co-expression of *wnt1* and *foxG* mainly in muscular and neuronal cells. **b** Tailless phenotype in *foxG* (RNAi) animals. **c** Heat map expression of *foxG* and *wnt1* in *wnt1* (RNAi) RNA-seq reveal that *foxG* is no down-regulated. Scale bar: 250 μm in a, 100 μm in b and c.

**Table S1.**

Specific accessible chromatin regions (ACR) of anterior and posterior wounds at 12 hR. For each ACR, it is shown the scaffold location, its position within, the fold change respect the opposite tissue and the FDR statistic value. We also show the specific enhancers of anterior and posterior wounds at 12 hR. For each specific anterior and posterior enhancer, it is shown the scaffold location, its position within and how it behaves in *notum* or *wnt1* (RNAi) conditions; being: accessible, slightly accessible, less accessible or non-accessible.

**Table S2.**

Results of the *wnt1* (RNAi) RNA-seq experiment. Dif Expressed is shown all the genes up- and down-regulated in the different regenerative time points. Down-regulated is shown just the genes down-regulated in the different regenerative time points including the presence of TCF motifs in the CREs (promoters and enhancers) and their presence in the dataset (46). Per genes is shown logFC, adj. P value, genome ID (39), condition (regenerative time point), gene name, Transcriptome ID (96) and its human homolog.

**Table S3.**

*wnt1* (RNAi) RNA-seq down-regulated genes with a TCF binding site in its promoter or enhancer region.

**Table S4.**

Presence of accessible chromatin regions (ACRs) with ATAC or DNasa evidences, and active enhancers with ChIP evidences in the first intron of *wnt1* gene in different species. – indicated no evidences. The genome source per each species was added.

**Supplemental Data 1**

Genomic sequence of *Smed-wnt1* locus showing the genomic organization of the gene (exons in green), the enhancers in the first intron (in yellow) and the FoxG binding motives (grey).

