## Supplemental Data 1 for "Genomic analyses reveal FoxG as an upstream regulator of *wnt1* required for posterior identity specification in planarians"

>dd_Smes_g4_197 dd_Smes_g4_197:263306..277425 (+ strand) class=gene length=14120

TTAATATATTTCAAATATCTTTATTATTAAATTTTAATTTGATTGGATAAAAATGAGGAGTTTAAACCAATTAAAATTCAAACTCTGAGAATAAAAGATGGAATAAATTACTTTAAATCATAACTACCTCAAAATCGAATTTTACACTCATTTATATTTTAAAATTTATTAACAGAAAATTTTTATTATTACTGATGAATCATTCCGGCGCTTCCATTTGTTTACCACTTATTATCAAAATCTTTTTGATAAGTTTTTCATTTTGTTTGACTTTGTCGATTAATAGAAATTGGAACAATCAAATATTCGTCAATCATCTCAGAAACACCAGATGGTGGTTAGTATTTATAATTTAAAATTTATAAATTATTTTCTGATATCCTTTGGTAATGAAGCAAATAATTATGTATATTTAATTTGAATATTTATGATATTTTTTGATAATATTAAAATTGTTTTAAAAATTGCTTTTATTATTATTATTATTATTATTATTATTATTATTATTGTTATTATTATTGTTATTATTATTATTATTATTATTATTATTATTATCATTATTATTATTATTGTTATTATTATTATTATTATTGTTATTATTATTATTATTATTATTATTATTATTATTATTATTATTATTATTATTATTATTATTGTTATTATTATTATTATTATTGTTGTTGTTATTATTATTATTATTATTATTATTATTATTATTATTATTATTATTATTATTATTATTATTATTATTATTATTATTATTATTATTATTATTATTATCTGCTTATTGTTAAAATTAAATATATTTTGTGATTGTTAAATGTAAACACTTTAAGAAAAATCTAAAATATTTTTACAAATTCTGTGACAAAATATTTTTTAGATATTTAATTATCACTATTTGATCTAATGTTTTTTAGATTAGTTATTAAAAATATTCATTATAATTAATACGGTCGGGAATAATTAATAAAAATTGTCTAAAAGATTGCAATATTTAAAATTAAATAAAAAATTACAAAAAAAAAATTATAATAAAATTTTTTCTCTTAAATTTCAAGAAAAATTGAAGTTTTCAAATCAGCGATGTCTGAAAATTTCGGCAATAATTTAAAAAGACAGTAATAAAATATTTGACACCAGCACCGTAAGCTAAAATTTTGCAAAATATCATTTTCTGTAATTATAAATAGTAAGTAATAAAAATAATTAGCATATGTATTAAGCAAGTTTGTCTTGAAAACAGTAAATGTATTTCATTGCCAATGTCAATGCGTGCAATAATAACTAATTTGGATTTATAACTTCGTACTGTAATTGTTGATTTTTCTTGTTTTTGTTTTTTTTTTCAGCAAAAAACATATTTTTCAAAATAATTCCAGAATTAATGTGGCGTTTAAAATTTTTTTTTATTTTTGGGCAATTAAAAATTTTTCCCATGAAATTTTGGTAGGCGGAAAAAATGATAAATGTGATTAATAAGTGACCTACCTCCGTTTAATGATGAGACTGACTACTCTAAAACGGTCTGATTGCCCACAACCACGTGGCCAATGTACACAAATGTGTCAATGACAGATATTGTCAATTTTGTGTGGCTTGAATTCAATCAAACATAAATCATAGCACTAGTAAACAGTTGATATAATCTATTTTCGAGTTTCTTTCAATTTGACCACCATAGTTTTGAAGAGTAATTGTTATGCTAAACCACACAGACACATGGATAAATAGGCGTTTTAATTGGGAGGGAGTAAAAGTAGCCCAATATTTCCCATGGCGAATATTATATCAAATTACTAAGCCAATTTGTCTAGTGTACACGAAATAGATTGAAGCATATTAATAATTACATTCTTCCTTCTCCTTCAACCGAAACCTTAACAATATTATTCCAATTCCTTAAACATAAACTTCCACTTTTTTTAATAATATTTTTTATTATTTTTTAAGTTTTAGATAATTATTATTTTAATATAGAATATAATTAAAATCATTTATTATTGATTATTTTAAATTTCTATTTATTTTTTAAACATTTAAAACAAAATATTTAAAACAAACAATTTATTATTAGCTATTAAAAAATGATTAAAAGAAATTTCATTATTTCTAATCGAAATAATACTTAATAATAATAATGGTTATTTAGTTTTCCATAGAAAAAAATTATATATTATTTTACTAAAAATTGTTAATGTTGAAAAATAGTAAAACTAAAATTGTTACATTCCAAAATCTATTAAAAGTTTTCTTTTTCATAAATGGAATTAATTTTAAAAATTTTTTAAATTCATAAAATAAAAATAAAACTACAATTTATCATTTTTTTTAAAACGATTAAATGATGTAAAAATAAATTATTTTTATAATAATTATATATTATTTTGCTAATGTTTGCAAATGTTTAAAAAAAGCTTAAAACTACAATTGTTACATTCCATTTTCCATTTAAATAATTCTTTATAAAGATAATTGTTTTTTAGAATTTTTTTCAAATATATACAATAAAAATAATGCAAAGATTTTTGTAAAAATAATTTAATTATATTTCTTAGTTAAATTAAATATCATTAGATCATCTAATATTATACTGTTGAAATTACGCGAGAGATGGAAATTCCTACACGGAGCTTGGATTAATGATTCCAAGTGCAGTTTTATTGCACTTCTAAAAAAGAAAATGTGTTTCAACAACTACCTAAATAAAAATAAATTTATAAAAAATATATATAGTTGACAGCTTTATTTCGCTTTCAAATTAACATTATCGTTTTCAAAAAAAATTTTTTTTGATAGATTAAAACAAAAAGAAATCATAATTAAGTCAAATGGGTATTTGCTAAAGGTTAATTAATCAGAGAAAAAAATATATAATTAAATAATAATTGATTAGGATAAAATTACTAAAAAATATTTTATACAACAACAACAACAACAACAAGAACAAGAACGGGTCAGTGTGAAATGATAAAATGAATGCACATCGACTGGATTTGATAGATGATTTGATTGGGGTTGGGAAAGCGAATTCTTTAAATGGATGTTGGCAATCAAAACGGGGCGGGCCTGAAATCGCCAAGGTCGAAATAACTGTCAATCACTGTGTAGTCAACATGACTGAGTTGAGACAGCTACCCGAGTAAATATTTACCTTAGTTACGAGTAAAATTCTTACTGATTACATCAGCTTAATCAACGTATTTTGCTGGGCCAGGCCTGCCAATAATCAAATATCAGTCTCTCCCAGCTGACAGATCCTATTTCCCTTATCAAGCACGTCAGTTAGTATTGAATTATAATTTATTGATAGAACAAATTAAAATATCACCTTTTAAAAATTAACATTCTCTTCCTCTTTCCAGATTAGTTGTACCAAATGATAATTTATTATTTATATATATATTTATATATAAGCCATTAATTTTGCGAAATGTAATGAAATTCGAATTATTCAACATTTTATAATAACAATTTTTTTATGAACTTTGCGTGGAAATTATTCTAAACTACAAAATTGGAAATTTATTTTAGTTTTAGTTTTTCTTCGTTTTAAATCTTTGATAATTAAAACAAATTCCATATTATCAATTTTTTTGAAGATTATAATTATAAAAATAAACACTCTCTTTTTCTTTCCAGAATTAAGTTGTTCCAAAAGATATTTTATTATTTATAATTTTTTTTTATAAGCCATTAATTTTGTGAAATGCAATGAAATTTGAATTATTTAACGTTTTATTATAACAACTTTTTTTATGAACTTTGCATGTAAACTATTCTAAACTATAAAATTAAAAAATTATTCTATTATAGTTTTTTTACATTTTAAACTTTGATAATTCAAAAAAATCCTATATATTGTCAATTTTTGAAATTAAAAACTTGTAATTATTTTTTTACGAATTTATGAATATCAATTCAGTTATTTAAACAAAGAAATATATATTTAAAAATATATATTTAAAACAAGTAAAAATATAAAGTTTTAAAAAGTAGCTTTTTCTTCCTCTTTTCAGATTCATTGTACCAAATGATAATTTATTATTTTTATAGTCATTAATTTTTATTAAATGTAATGAAATCGAATTATTTAACAATTTTTAGTAACAATTTTTTACATTTAAAATTTTGATAATTCAAACAAATCCTATATTGTCAATTTTTGAAAATAAAAACTTGTAATTATTTTAACGAATTTATGAATATTAACTTAGTTATTTAAACAAAGAAATACTGATTTATAACTATATTTATAAAACAAATTAAAATATCAACTTTTAAAAAGTAGCTTTTTATTCCTCTTTTCAAATTCGTTGTACCAAATGATAATTTATTATTTATATAGTCATTAATTTTTATTAAATGTAATGAAATCGAATTATTTAACATTTTTTAGTAACAATTTTTTATGAACATTGTGTGTTAATTATTTTATTCTATAAATCGAGAATTTATTTTACTTTATTAAAACAAAAATATTTTGAATTTTAATTAAACTCTTTTATTTATTAATATAACAATTTAAAATATCACCTATAATTTCTTATTTAAATTTTCTATAAGCCAAAAATTTTGTAATACAATTGAACTATGTAAAATTTAATAATTCTGTAATAAAAATGTTTTGTGAAATTTGCTTGTTAGTTATTTTAAACAACAAATCGGAAATTTATTTTACTTTTATTTTTATCATTAAATTATTATTTTTTTGATAAATTATGATAAAACTTATACAGTGATTGTTGTTCGAATAAAGCACTTGTTTAAAATGTCATAATTAAAACAAATTTCATATAATCTGTTTATGAAAGTCATAATTTATTGATAGAACAAATTTAAAACACCATCTTTTAAAAGTGAATATTTTTTTAATCTTCCAAAATTAATTGTAATATATATAATAACTTATTATTTAAAATTTTTTAGACTATTAATTTAGTGAAATGTAATGAAATTGAATTATTACATTTAATATTATGCAAGTAGAATTTTTTGGAAACTTTGCGTGTTAATAATTTTAAACTACAAAATTGAAAATTCATTTTTTATGTGACAGCAATTTAATCATGAAAATGCTTTTTTTGTAAAATATGTACAGCGATAGTTGTTTTAATAAAGTTCATGTCTAAAATTTGATAATTAAAACAAATTCTATATGATCAATTTTAAAAATTAAATTATTTTTTTATTTCCTTTTATGCTTATTTTAATTAGTTACTAAAACAATATTAAATTACAGCATAATAAAATATATTTTATTAAGTATAGAAAATTTTAAAACAAGTTTGAACAGAGTCGATTTTTACACTCACTTTTTCCCCTGCAATATATTTGTACATCATAATGCTAAATTTATAAAAATTTATATTTTATTCTGCATAAGTTTAAAATCAAAAAATGTATCGTTGTCCCTCTTAGTCATCTGTATTATTTTGGTTGCTCTGCTTTTTGTGGAAATGTACTTCAGTGGTATATTTTAAACCTTTTACCTATTAACCCCCCTTACTTGCACATTCAAACAATAAACCTCTTTTCCATATATATACTTATTTTTTAAATTAAAATTTGGATAATTAAATATCAATTAAAAATAATAAACAAATACTAACATATAAAAAATGGTCAAAATTACTTTTACAATAAAAATAATGCAAAAACGAATATTAGTAGCTTGTAGTTTTTCATAATTTATTTTTAAGTAAATATTTCGACGCGCAGTGAAAATTTAAAAAGTAATTGAAATATTTTTAATCTAGGCATTTATATAAAATGAAAAGTGATCCAAATATTGCAACTAACAGCCTTCAATATGCTTTCACCAGACACCAATGGAATCAATTTTACAGTTTTGATCCAATTAGACGTAAGCTTTATTCTCAAGAGTACAGACCCAATTGGAAAAACTTTACCGATTTGCCTGCTAACAGTGATCTAATTCAAGTGGCTATTGAAGGAATCAGAAAGGGTATTTACACATGTCAAAAGTTGTTTGCCAATCATCGCTGGAACTGTCCGACACCGAATTTGAATAATCCATCCGCCTTACTTTTTGGTGACATTATGCTCAAAGGTAATCAAATGAAGTTTTTTTTTGAAAAATAACTTAAATTTATTATTTTTAAGGATTTCCCGAAACCGCTTTCATTTATGCGATGCTTAGTGCGAGTGTCGCTCAAACTGTCGCTGAGGCTTGTTCGTTTAAGTTATCACACTGTCCGTGCAATAATAAAGGACGGATATCTCAAACCAACTGGATTTGGCAAGGTAAATTGTAATTATTTATTTTTAGAAAGATTGCACAAATAATAATAATAATAATAATAATAATAATAATAATAATAATAATAATAATAATAATAATAATAATAATAATAATAATAATAATAATAATAATAATAATAATAATAATAATAATAATAATAATAATAATAATAATAATAATAATAATAATAATAATAATAATAATAATAATAATAATAATAACAATAATAATAATAATAATAATAATAATAATAAATATAATAATAATTAATAATAATAATAATAATAAAAGCTTTATAAATATGTAATTTAATATGTGTTTTGTAAGCTATTACTGATGAAATAGATACTGAAAACGTTGAATTTATTTAGGATGTGACGACAACGTACAATTCGGTCGAAGATTTGCGAGAAAAATGTTTGATCCAGACATTCCAGATTGGAATAAAAAGACATTAATGAACCTTCACAACACGGGAACGGGTAGAAGAGTAGGTTGAGATTTTTTAGTATAAACATGAGTTTAAGATACAATTTAATTTTTTCACAAACAAACAATTTTTTTTTATTTTTAATTTCCTTTTATATTCCAATCATACAGACTTATCAATAATCGGCCTGTTGGAAAACAAATTTCGGCAAATGCCAATTTTCTTAAAAAGTACGCTAATCCGAAAGAAAAGCACCTTTAAACATAATTAAACAAAGTAAATTCTCTTTTTATATTATTTAAACATATTAAGGAGGAAAACGGTCTAGTATATGCTTTTAAAATCAATAAAATTTATACTTTTTCAAAACAGACTGGAAAAAAAATTTCGGCAATTATATTTAATAAGGTATATGTTTCCCTTTGGCTCGAATCACACAATTAACTCTATTGTTCATAGTCTTTACGAGTCTTTTACAAAGCTCTGAATCGATATTTTCCCATATTTTTATAACATACTGCTCAAGTTCTCTTAAATTAGTCGGCTTTCTTTTCCTTACTTCTCTCTTCACGTAATTCCATAAATGTTCAATGGGATTCAAATTTGGAGATTGTGCTGGCAAATCTAATAAGTATTCGCCTACAACTCTGGAAGTATGTTTAGGATCATTATCCTGCTGAAAAACAAATTCATCATCGAGACTCATATCTCTTGCCGAAGGACCTAGATTTTTAGTTAGAATGGACTTATACATTAGACCATCCATCTTTCCCTTGATAATATGGAGTCTCCCGACACCTTTAGATGATATGCATCCCCATAGCATTATACTACCCCCTCCATGCTTTATAGTTGGAAGACAATTTTTAGATTCTAAGGCCATTCCTTTCTTTCTATAGAATCTCTGCCGCCCATCGGATGAAAAAAGGTTAATCTTTGTTTCATCGCTCCAAATTAACTTATCCCAATCACTTTCTTCCCAAGAAGACCATTCATTTGCCAAATCCAATCTTTTCATAATATTTGAATTGGTAAATTTCGGTACCTTCCTTGGGACTCGAGATCTATAAGAGGTATCTTTGATATAGTTTCTTACTGTTCTGCTTGTAACGGATTTTTTTAGTAGTCTTTTCTATTTCATTTTTGATACTCTAAGATGACGTAAACGGATTTTTTTCAATTTCATTTAAAATAATCTGGATATCTGAATCATCCAATGACTTTTTTCTTCCAGAACCAGACAATCTCAATACAGTATCATGTTTATCATATTTCTTTATGATACTGTATATTGTAGACCTTGGAATATCCAGTCTTTCGGAAATTTCAGAATCCGAGAGGTTTCCTTGATGCAACATTACTACCTCTGAATCACGTACAATTTTAGTTTCATTTATGCTCATTAAAGGGTAATAAATTATTCCACAATTTTATTTTTTGGCGGGAAAAGTTTTTGAGAGGTTTGTAGAAATTTGCCGATTTTTATTTTCCAGGCATATAATAAAATTTTTAGATTTAAACAGAAAATCAATTATAATATCGAAAATAAAAAGCATAAACCATGTTGAACTTGTTAGTTTTAATACATGCCAAATTTTCATTAATAAATAATGATATTATTAAGGACTAATTCGAAAAAAAGACATTTGCCGAAATTTTTTTTCCAACAGTGTATATATATATATATATATTCGCCATTAAACGTGCCATTAGAAGTGCCGTTAAATTGTACATAACCAATAGGCGGCCTCTGTGTCTCATAAGAACAATTAATATAGTTTTATCATATATAACTTACGGATTTATCTGGGAATACCTATGTACTCGTTTAGACTAATCTGTATTTCTGAAAATCTAATAATCAAATTTCCACATGTTCTTTGGAAATTTTTAAAATCATGTTTTGACCCGAAATTGTGGCAATTTTCAATATTTGAGTAAAAATAATAATATAAATAAAAAATGAGAGGAATTTTATTTTTGTGATTTGGAATTCAATTATTCTTTTTTACTATTATACTTATATAGGTTGCAGTCAAAAGTATGAACGTGAAATGTGTATGCCAGGGCACAAGTGGATCTTGTACGACCAAAATTTGCCACAGAAAGGTTGCTAGTATTGACGAAATCGGTAATATTCTTAAAGAAAAATATGAAAATGCCGTAAAGGTCGTCAAAGGTAACAAGTTATTTTATTGAAATTATATACTGAATAAATGTGAAATACTTAAACCAAAAGTTATAAATATGTTTAAAATATTGTTGAAAATTTTGATAAAATATTACTGAGTTGATTAAGAATAATAGAAGATTTATTGGAAATTTACAAAAGTTCAATGAAAGCATTTTTTGTAAAAATTAATCATAATTTATACAAATAAATAATTAATTAAAAATGAATTGTCTAGAATGGAGCTGCCATGAAACCAAAATTTATTTTTATAATAAATTAAAAATTTTTAAATTCTATACTTAAATTGTTTATCTGCTTACTGTAGAAATCAAGATGTTTTAATAACTATGGGATTATAAAATGAATCACAGAATGGAATTATTCAAACAAAATAGTGTAATTTATGTTGAATGAATGATCATAGGTGAACTTGTTTTTCTCGATTTGAATCTCAACAGTTTTTTGTAACTCGAAAAATTGTTTTACTAAAGTAAATTGTTTACGCATATCTGATTACAGAAAAGATTTGGTTGTTCTGACGATTAAAATAACTCGCGCAAATATTAAACAAACAACGCACTGCATTCATTACCATATAATGCTATAGAAGAGTAAAGTTTTTGATTTTTTGATTTTCTTATTGAAAATCAAATTTGTTTATTTTAAATAACAATTTAAGTTACAGGTTAAATAAGAATATCGTCATTATTTAGAATAAATGTTTCCACGTAAATAAATAACAGCAATTTTCAACAAGTTCTTATATCAAAGGCAGTTTCTGTAAATAATTGTTTACAAGTTCCATTTGATTCTTTAATCACTTTTCAAATTGCCAACAAAAACAAAATTTAAAAGTTATTTTTCTTTTGCGTATTTAATGTCTAAAAAACCCCTTCATAAGTCCTTCCGATCTTGAACGTTAAATTGATTCAACCAATAAAACGGATGCGCGAAATTTCGCTATTCAGCGAATGTAGTAAAATGTAAATAATTTGTTTTAATAATTTAACTGGATATAATTATTAATTTTTATTCCATTTAATCATTTTTTCTTTGGCATAAAATCTGTTTTTTTTAATTTAGACTTAATAATATTTTCTTTGTTTGTATGGAATGCCAAATTTTATTTAAAAGTAATTTAATAAAATTTTACAATTTTCAGTTGACATAATTCATGGAAAATATTTCGTATATTAGGAGGTCTTAATAAATTAAACTCATCAACAATTTCTAAATTTTATGTAAATCTTTGCTGTAACATTTTCCAATAATTGGAAAAAGGCGGTTTGTAACTTATTTTACTGTGATAATGTAAATTACTTAAAATTACATCGAGTTTCAAATACCGAATTCGACAGTTTCGAGTTGGCGCCAATTCAATTAGAATTTTAGCAATGAAATTGAAACTTATTGATAAATTTAATAGAATTTTTAGATTAAATGATTTAATTATATAAAATTTCAAGAATTCTCATTCAATTGCTTTTTTATCAGATAATAATTCAGACAATTTTTGTAACGATTCAAACAAAATTTTTATTACAGAAATAAATCTAAACTTTTCCATGAAATATTTATAGTGATAAACAGAAAATTAAGAAAAAAGCTTTTAATATAATTTAGGTAAATTGCACTTCGTGGTATTAGGACTTTTAATTATGGGCAATAAATTATTTAAAATGAATTAAACTTAAGTTAAAACGAATTTAACAAGTAAGCCCTTTTAATTATCAAATTAGTAATCTCTTTAAAAGGGTGAAAAATGATGAACGAATACCAAAAGATTTTTAGGGGGTAAAATTTGTGTGTAATTACATAAATGTATATAATTATGTATTTATTATTAATCAATTTGGCAAAGAAGATTATATATATATACATATATATACTAATGCAATTTTTTGTTTTAATGATTGATTTGGTTGAATATCATTTAAACAAAGTTTTAATCAGCCGAGATCTGGGCAAAACAACGCATTGAAAACTGTTTGTACATTTATTCCAATTCCTTACTTAAATAAAACAAGTTTATAATTATAAAGTTTATTTAACAAATATAATATTTTTGAAAGTAGAAAATATAAGCTTGTAGTTAGCAGCATAAAATAAATGTTAAACAAATTTTTAATATTTTTTAATTTAATTATTAAAAATGTAATTCTACTAAATCGCTAATATAATGTTAAACCTTGATCATATGAAACAATGTCAATTCAGTTTTGTTGCACAATCTGTAAGTCTTGGGTATAGATTTTACATAAACATATTTATGTAAACTAAATAAATTATTTATTCCTGCACCGCCAATAGCTCAAGTAAATAATAAACAACATTTTTATAATCTGTTTTAAGTTATTTCAAGAAATTATAGAAATGAACGACGTTTCGCATTGTTTACTTCAGAAATTCCAATATTCCCTGGAAATTTCATTGAAGAAAATATTTAGCTGTATAATGTTTCATGACAGATGATACTGATCACATAAATCAACCCTAAATGCTTCTATGTTGGCAAGAGGAAGTCTTCTTCCATTCTGGTACTCTCAAAATAGACACTTGATAAACATCATATTGTTAATTAAAGTTGTTGAATGTTAATTATAACGTTAAATAAAATGAAATATGTATGGTAAATAATTAATGTATTAAATTAATTTTCTGTTAAATAATTAATTTATTAAATAATTAACTTTGTTTTATTATGAATAATAATAATAACAATAATAATAATAATAATAATTATAATTATAATAATAATAATAATAATAATAATTATAATAATAATTATAATAATAATTATAATAATAATAATAATAATAATAATAATAATAATAATTATAATAATAATAATAATAATTATTATTATTATTATTATTATTATTATATTAAAAGTTTGTGGCAAATACTTATAATTAGTTCTAATAATGAAATTGAAAGTCCTTAAGAATACAAGTCCAGGTAGTTGTTTGCCAACTTCAGATATACCACATAATTCAATTATATATTATCCACAAATATATTCAAATTCGACGTTACAAAAATCATAAATCATGTTTTTATTTATATTTCTAATTTCATTTGAATGGCAGAATAACAAATGAATGTCATTTGGTAACCCCCAGTTTCCTTTGCAAAATCTCCCAATCTATAGGTCAAGTGATAGAGTAAGTAATTCTGAAATAATTTTTGTTTCAGCTTAATTCCAAAATTATAAAAACGATGCTCAGAAATTTATTTATTTATTGTTTTCATTTATTTCAATATACAATAGAGAAAAAAATTAGTGAAATACAAAAATATAATTAAATGAAATATGAACTCATTGGTAAAATTTTATTAAGACAAATATTAAAACAAAAATATATTTGGGAAAATAAAATAAATTTAAAACGGCTACTTTAATTGAAATATTTTTTAATTGATCAAAATTTGATTACGTAAGGCTGTCTGTCTGGTGCAAAATGTTGACATCTTTATTAACAAATTAAAGCAGATGTAGCCTGCAAAAATAAAGAATGAATTTAATACACTGATTATTTTAACTGAAGTCATAAAAGTTGAATAGCTTCAAAAATGTGGTCTTGCTGCCTTTTGGTGAATCAATTTAACTATTTCTGGTCAATAATAACTGTGCTTGTTTATGTTTTATTTATGACTTACGCAGACTGTGTTTGACTTAATTACCCAGAAATTCTCAAAAAATTATAAATTAATCTTTTACTAGCTCATGTTTTTTTGGAAATAGTCAACACTTGACAGCCAAATATATTTTTTGAAAATTAAAATACAACAAAAAATTTTAAAAAAACTAAAAATATTCATTTAAAGTTGTAAAATAAATAACCAAAATGAAATAATCTAAACCCAGAAACAATTTTTGATTACAAATATTTTATAAAGTTAAAAATGTTTTTATTATTATTATTATTATTATTATATCAGATAAGTGAATAACAAAATAAGCCCTTGCTACGAAGTAATAATCGAAACTTTTTTCAAAAGTATAATTATTACTTCGTAGCAAGGGCTTATTTTGTTATTCACTTAGCTGTTATATTAATTTAGGCTGTTACCAAGGAATCATCGTTTTATACTTATACTATTATTATTATTATTATTATTATTATTATTATTATTATTATTATTATTATTATTATTATTATTATTATTATTATTATTATTTTTATTATTATTATTATGATGATGATGATGATGATGATGATGATGATGATGATGATGATAATGATAATGATAATGATGATGATGATGATGATGATGATGATGATAAAAATAATAATAATAATAATAATAATGATAATCAAAAAAAATTGGTAAAAGATATGATAAGCCTTTGCTGCTAGTTTCAAGTTATTTAAAAAAAATGATGAGAATTTTTATGAAATTGCGTTATCTTATTGAGAGATTGAAGTAAACTATTTCTTAGTCAACAAATCTTGTTATTTGTTTTAATAAATTTTACACGTGCGGCAAAGGTTTCTTGTATCCTTGGCTCATATTTTTGTAAATTGATGCTGTTACTATTAAATAATAGTTTATTACTGATAATAATAATAATAATAATAATAATAATAATAATAATAATAATAATAATAATAATAATAATAATAATAATAATAATAATAATAATAATAATAATAATAATAATGATAATAATAATAAAATTATTATATATTTTTAGAAGATAGTGAAGTTAACTTACTTGCTAATTCGCAGAAAAGAAAAACGACTAATAAAAACCAACAATATCGTAAGGCACTCATAATTGCAAGGAAACATATTTTCATTCGAACAGAAAAACAATATCCATATGCTAGAGATTTAGTTTATTTAGAAGATGTTCCAAAAAACTTTTATTGTGATTCAAAACCCGAATGGCACATTCTGGGCACGACGGGGAGAGTTTGCAATTCTTTGTCAAATTCGACTGACAGTTGCAATAATCTTTGCTGCAACCGACCTTTCCTAACAAGATCTAAAACAATAATGGAAAATTGTAATTGCAAATTCATTTGGTGTTGCAAAGTTGAATGTCAACAGTGTCAAAAAAGAATTTTTATTGAAACATGCCAATAATAATAATAACAAGAAAAACAAATCGTAACAAACGTCACAGTCAAATTTTATTATAGAAAATTATTTTCATTTAAATTCTTTTATTTAGAGTTGTGGAATTTTATTTTTGTCCCAAAGAAATTTATAAATTCAGCAACAATTAAATTTGCTTGTTTTAT

CDS wnt1

Enhancer wnt1

foxG motif
